# Discovery and Structure–Activity Relationship Optimization of a Novel Rv1625c Agonist Chemotype with Antitubercular Activity

**DOI:** 10.64898/2026.09.17.752203

**Authors:** Han Xie, Paridhi Sukheja, Kirsten Tolentino, Jasmine Webb, Ashley Woods, Victor Chi, Dipak Kathayat, Christine R. Montague, Brian C. VanderVen, Leonard Winneroski, Greg Durst, David Mendel, Philip Hipskind, Kenn Henry, Case W. McNamara, Baiyuan Yang, Arnab K. Chatterjee

## Abstract

Rv1625c/Cya has emerged as a promising target for the development of treatment-shortening therapies for tuberculosis. Screening of an Enamine compound library identified **sBQQ004** as an initial hit, and rapid hit optimization led to compound **1**, which was subsequently confirmed as an Rv1625c/Cya agonist. Structure–activity relationship studies identified lead compound **25** with potent activity against *Mycobacterium tuberculosis* H37Rv under cholesterol-dependent growth conditions (MIC = 0.27 µM) and strong intramacrophage activity (EC_50_ = 0.079 µM). Compounds **1** and **25** showed oral bioavailabilities of 84.5% and 51.9% in mice, respectively. Repeat BID dosing of compound **1** resulted in a dose- and time-dependent decrease in systemic exposure. Despite this pharmacokinetic limitation, the potency and overall profile of this chemotype encouraged continued optimization.

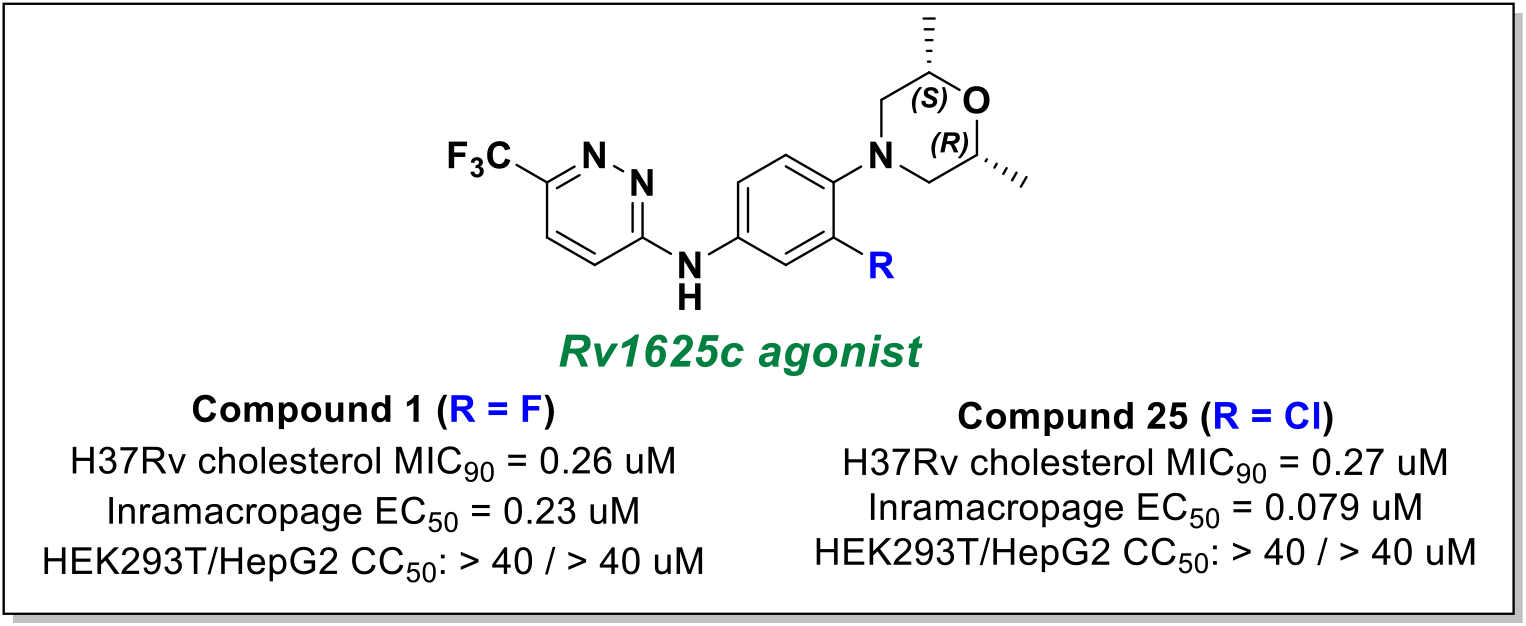

## Introduction

Tuberculosis (TB), caused by *mycobacterium tuberculosis* (Mtb), remains a major global health threat and is the leading cause of death from a single infectious agent. Despite substantial advances in TB prevention, diagnosis, and treatment, an estimated 10.7 million people developed TB and approximately 1.25 million died from the disease in 2023, imposing a substantial health and socioeconomic burden, particularly in low- and middle-income countries.^1,2^ Even for drug-susceptible TB, current treatment typically requires 4–6 months of multidrug therapy, creating challenges for patient adherence and treatment completion.^3^ In addition, the emergence and spread of drug-resistant Mtb strains further complicate treatment.^4^ Therefore, new anti-TB agents with novel mechanisms of action that can shorten treatment duration remain urgently needed.^5–7^

One promising strategy for identifying new therapeutic targets is to exploit metabolic pathways that are critical for Mtb survival and persistence within the host. Mtb occupies diverse intracellular and extracellular niches during infection, and primarily resides in macrophages and tissue lesions such as granulomas.^8^ Mtb can utilize host-derived lipids, including cholesterol, as carbon and energy source, to support its survival in lipid-rich environments as found in necrotic granulomas.^9^ Genetic studies have further demonstrated that disruption of pathways involved in gluconeogenesis, cholesterol utilization, or the methylcitrate cycle (MCC) can attenuate Mtb survival and virulence during infection. Within this context, Rv1625c (Cya), a membrane-bound adenylyl cyclase whose pharmacological activation perturbs cholesterol utilization in Mtb, has emerged as a promising therapeutic target.^10–13^ Rv1625c agonists, including mCLB073 (TBD11) and GSK2556286, have demonstrated potent anti-Mtb activity *in vitro* and efficacy *in vivo*,^14–15^ and have been advanced to clinical trials. Activation of Rv1625c elevates intracellular cAMP levels and disrupts cholesterol catabolism in Mtb, resulting in inhibition of bacterial growth, although the detailed downstream mechanism remains incompletely understood. More importantly, preclinical studies have demonstrated the potential of GSK2556286 to enhance the efficacy of BPa/BPaL-based regimens and so contribute to treatment shortening.^15^ Nevertheless, the identification of structurally distinct Rv1625c agonist remains an attractive strategy to mitigate clinical attrition risk, address scaffold-specific liabilities, including off-target toxicities and suboptimal pharmacokinetic properties, and reduce the potential impact of resistance that may emerge against existing chemotypes. Herein, we report the discovery and characterization of a novel chemical scaffold targeting Rv1625c. Detailed structure– activity relationship (SAR) studies identified analog **25** with improved anti-*M. tuberculosis* activity, and the pharmacokinetic properties of this chemotype were further characterized in mice.

## Results and discussion

### Identification and Initial Optimization of a Hit from Library Screening

A screening library comprising approximately 150,000 structurally diverse compounds from Enamine was evaluated against Mtb H37Rv in cholesterol-supplemented 7H12 medium using a 384-well format at a single concentration of 10 µM. Primary hits were defined as compounds exhibiting at least 90% inhibition of Mtb growth and were subsequently evaluated in dose-response assays to determine the concentration required for 90% growth inhibition (MIC_90_). In parallel, cytotoxicity (CC_50_) was assessed in in the mammalian cell lines HEK293T and HepG2. Hits were prioritized based on potency (MIC_90_ < 5 µM) and an acceptable selectivity index (SI = CC_50_/MIC_90_; SI > 10), then were evaluated in an intramacrophage assay to confirm activity against intracellular Mtb.

**sBVQ004** was identified as one of the prioritized hits, with an MIC_90_ of 0.74 µM in cholesterol medium, an intramacrophage THP-1 EC_50_ of 6.69 µM, an MIC_90_ of 3.7 µM in OADC medium, and no detectible cytotoxicity in HEK293T or HepG2 cells (CC_50_ > 40 uM). Notably, **sBVQ004** retained comparable activity against an Rv1625c-resistant Mtb strain in cholesterol medium, suggesting that its antibacterial activity was independent of Rv1625c (**Table 1**). To address the potential metabolic liability associated with the ester moiety of **sBVQ004**, Compound **1** was designed by replacing the ester with a trifluoromethyl group and introducing defined stereochemistry at the dimethyl-substituted morpholine ring, in contrast to the racemic **sBVQ004**. Compound **1** improved both activity in cholesterol medium (MIC_90_ = 0.26 µM, intramacrophage activity (EC_50_ = 0.23 µM), and to our surprise it lost activity in OADC medium (MIC_90_ > 40 µM), resulting in a conditional activity profile resembling that of the previously characterized Rv1625c agonist mCLB073. Consistent with this hypothesis, compound **1** lost activity against an Rv1625c-resistant Mtb strain, indicating that its antibacterial activity is dependent on functional Rv1625c and suggesting Rv1625c as its putative target. To further investigate its mechanism of action, compound **1** was evaluated using a cAMP-responsive Mtb reporter strain (Gflamp-1) (**Figure 1A**). Mtb-Gflamp-1 constitutively expresses a codon optimized Gflamp-1 and mCherry allowing flow cytometric quantification of cAMP levels in the bacteria. Gflamp-1 is a circularly permuted green fluorescent protein that carries a bacterial cAMP binding domain to produce a GFP signal when cAMP levels increase in the bacteria.^17^ Treatment with compound **1** increased GFP reporter activity relative to the DMSO control and produced a response similar to that of mCLB073, consistent with activation of Rv1625c-mediated cAMP signaling. We further evaluated compound **1** in the Hu-Coates model against slowly replicating and non-replicating Mtb (**Figure 1B**).^18^ Addition of compound **1** to the three-drug combination of bedaquiline (B), pretomanid (Pa), and linezolid (L) (BPaL) further enhanced antibacterial activity, reducing bacterial burden by about 1.3 log_10_ CFU compared with BPaL treatment alone. This effect recapitulated that observed with mCLB073. Taken together with the Rv1625c-dependent resistance and cAMP reporter results described above, these findings support compound **1** as an Rv1625c agonist.

**Table 1.**
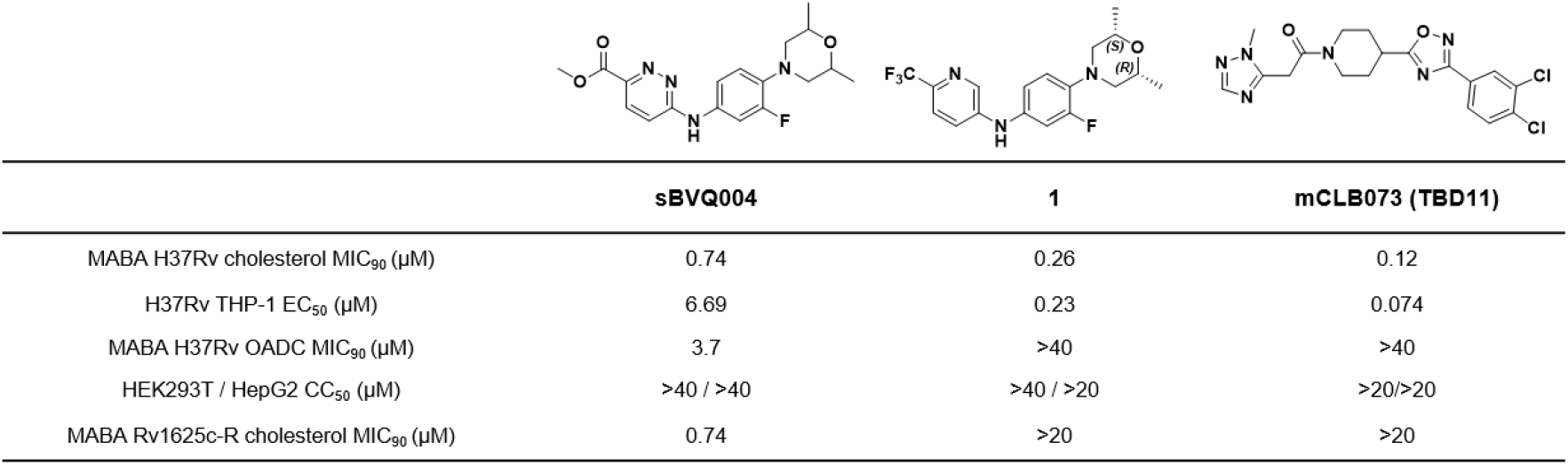
Structures and biological profiling of hit compound **sBVQ004**, initial lead 1 and known Rv1625c agonist mCLB073 (TBD11)

|  |  |  |  |
| --- | --- | --- | --- |
|  | **sBVQ004** | **1** | **mCLB073 (TBD11)** |
| MABA H37Rv cholesterol MIC<sub>90</sub> (μM) | 0.74 | 0.26 | 0.12 |
| H37Rv THP-1 EC<sub>50</sub> (μM) | 6.69 | 0.23 | 0.074 |
| MABA H37Rv OADC MIC<sub>90</sub> (μM) | 3.7 | >40 | >40 |
| HEK293T / HepG2 CC<sub>50</sub> (μM) | >40 / >40 | >40 / >20 | >20 / >20 |
| MABA Rv1625c-R cholesterol MIC<sub>90</sub> (μM) | 0.74 | >20 | >20 |

**Figure 1.**
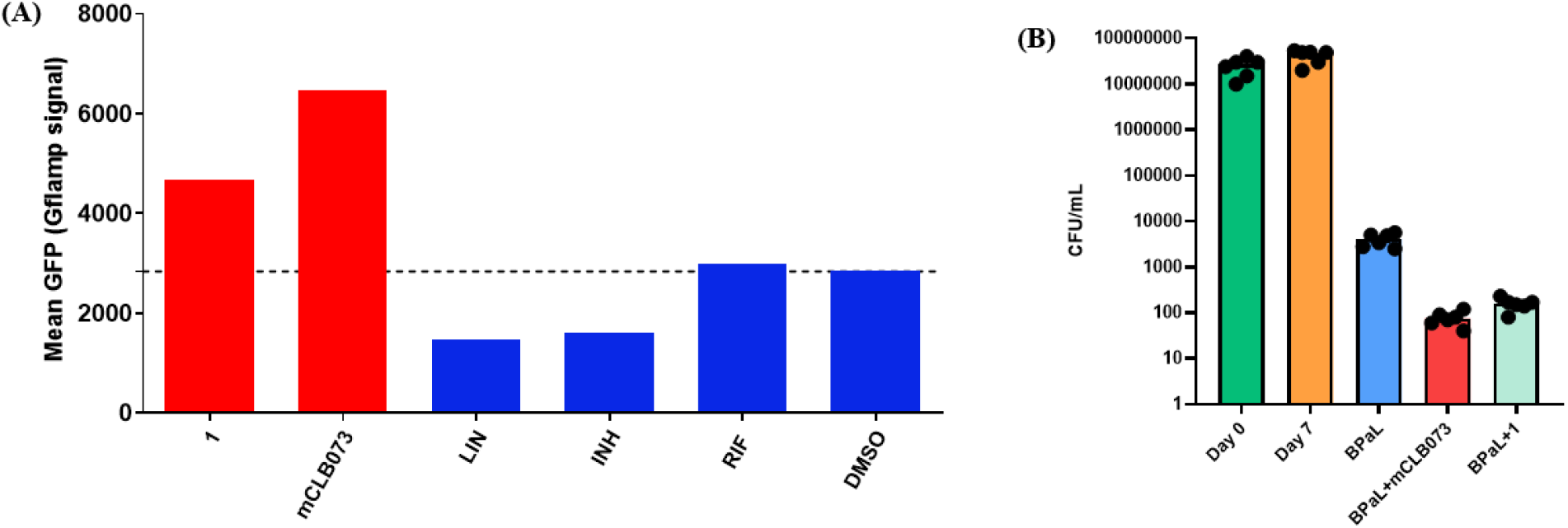
**A.** Rv1625c/Cya activation measured with the cAMP responsive Mtb Gflamp-1 reporter strain. The mean fluorescent intensity from ~20,000 bacterial cells is depicted; **Figure 1B**. Bacterial activity of compound **1** and **mCLB073** in combination with BPaL in Hu-Coates nonreplicating persistence (NRP) model.

**Figure 2.**
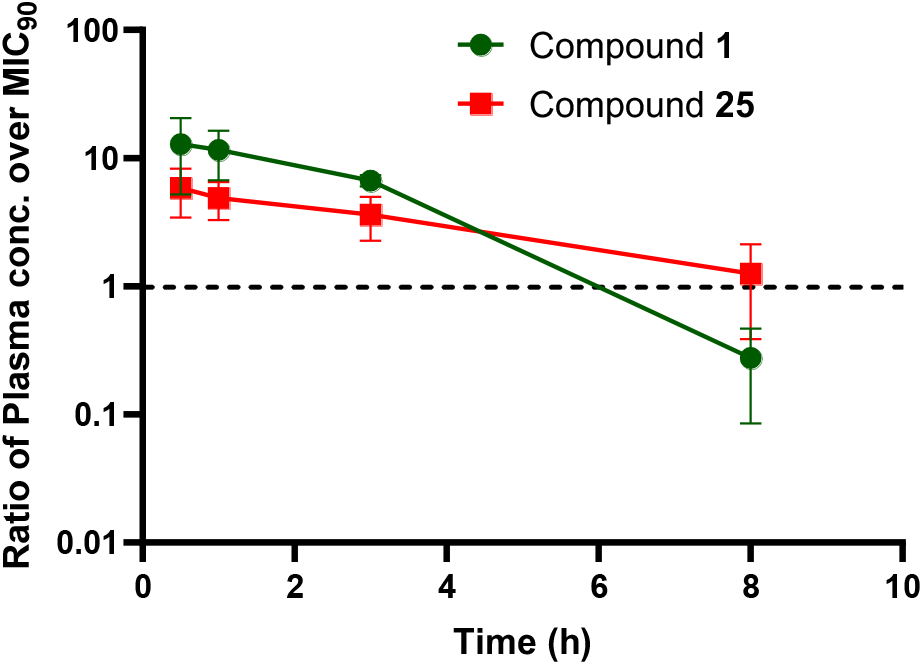
Ratio of mean mouse plasma concentration to the corresponding cholesterol MIC_90_ for compounds **1** (green) and **25** (red) following a single oral dose of 20 mg/kg in a solution formulation of 75% PEG/25% D5W

### Structure–activity relationship (SAR) optimization of compound 1

With compound **1** identified as an Rv1625c agonist, we next undertook a systematic structure–activity relationship (SAR) study to improve antimycobacterial activity and physicochemical properties. Compounds were evaluated for activity against *M. tuberculosis* in both a cholesterol-containing growth assay and an intramacrophage assay, together with cytotoxicity assessment in HEK293T and HepG2 cells. The scaffold was divided into three regions for optimization: the left-hand heterocycle, the fluoro-substituted phenyl core, and the right-hand morpholine moiety. The target compounds were prepared according to the route described in **Scheme 1**. The synthesis started with an *S*_*N*_Ar reaction between 1,2-difluoro-4-nitrobenzene (**S1**) and (*2R,6S*)-2,6-dimethylmorpholine (**S2**) to afford intermediate **S3**. Reduction of the nitro group of **S3** with Fe/NH_4_Cl provided the corresponding aniline intermediate **S4**. Subsequent coupling of **S4** with the chlorinated heteroaryl under PTSA (*p*-toluenesulfonic acid)-mediated conditions provided compound **1**.

**Scheme 1.**
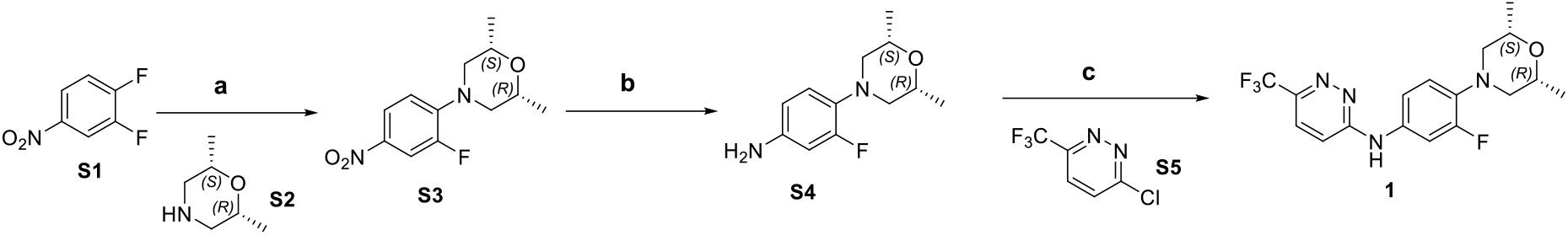
General Synthesis of Target compound **1** and analogs Reagents and conditions: a) (2R,6S)-2,6-dimethylmorpholine (**S2**), K_2_CO_3_, DMF, rt, 16h; b) Fe, NH_4_Cl, EtOH/THF/H_2_O, 80 °C, 4h; c) 3-chloro-6-(trifluoromethyl)pyridazine (**S5**), PTSA,1,4-dioxane, 110 °C, 4h

We initially focused on modification of the heteroaryl region because of its synthetic accessibility (**Table 2**). Replacement of the trifluoromethyl group with weak electron-withdrawing substituents, including a 1,2,4-oxadiazole (**7**) and an amide (**8**), or with electron-donating alkyl groups (**9** and **11**), resulted in substantial losses in activity, with at least a 10-fold reduction in potency relative to compound **1**. In contrast, strongly electron-withdrawing substituents, including cyano (**3**) and difluoromethyl (**2**), were better tolerated, showing less than a 5-fold loss of potency in the intramacrophage assay, although larger reductions in activity were observed in the cholesterol-containing assay. These results suggest that the trifluoromethyl substitution makes an important contribution to the antimycobacterial activity of this series. We also investigated replacement of the pyridazine ring with alternative heterocycles. Analogs containing pyrazine (**4**) or pyrimidine (**6** and **10**) rings were inactive in both the intramacrophage and cholesterol-containing assays. The 1,3,4-thiadiazole analog **5** retained good activity in cholesterol-containing medium (MIC_90_ = 0.66 µM) but exhibited decreased activity in the intramacrophage assay (EC_50_ = 3.48 uM), making it less attractive compared to compound **1**. Most analogs showed little or no cytotoxicity against HEK293T and HepG2 cells, with only compound 4 exhibiting moderate inhibition.

**Table 2.** *In vitro* anti-TB inhibitory activity and cytotoxicity of the heteroaryl analogs 1-11.

| Compound | R<sub>1</sub> | TB activities |  | Cytotoxicity |  |
| --- | --- | --- | --- | --- | --- |
|  |  | Intramacrophage<br>H37Rv THP-1<br>EC<sub>50</sub> (uM) | H37Rv Cholesterol<br>MIC<sub>90</sub> (uM) | HEK293T<br>CC<sub>50</sub> (uM) | HEPG2<br>CC<sub>50</sub> (uM) |
| 1 |  | 0.23 ± 0.15 | 0.26 ± 0.17 | > 20 | > 40 |
| 2 |  | 0.62 ± 0.32 | 1.33 ± 1.28 | >40 | >40 |
| 3 |  | 0.87 ± 0.41 | 2.43 ± 0.34 | >40 | >40 |
| 4 |  | 3.97 ± 1.19 | nd | 18.45 | 25.89 |
| 5 |  | 3.48 ± 3.25 | 0.66 ± 0.34 | nd | 37.17 |
| 6 |  | >20 | 8.16 ± 3.79 | nd | >80 |
| 7 |  | 19.25 ± 7.09 | nd | >40 | >40 |
| 8 |  | >20 | >20 | >40 | >40 |
| 9 |  | >20 | 10.16 ± 7.15 | >40 | >40 |
| 10 |  | >20 | >25 | nd | >80 |
| 11 |  | >20 | >20 | >40 | >40 |

nd: not determined

Next, the morpholine domain SAR was investigated (**Table 3**). The two *trans* stereoisomers **13** and **14** showed more than a 20-fold reduction in potency compared with the meso *cis* isomer **1**, indicating a strong stereochemical preference for the cis configuration. Ring opening (**12**), introduction of additional methyl substituents (**16**), and incorporation of bridged ether motifs (**18** and **19**) abolished activity in both the intramacrophage and cholesterol-containing assays. Interestingly, dimethyl piperidine analog **15**, retained comparable MIC activity in the cholesterol-containing assay but showed an approximately 50-fold reduction in potency in the intramacrophage assay. Introduction of a polar sulfonyl group (**17**) also resulted in complete loss of activity. The mono-trifluoromethyl-substituted analogs **20** and **21** are tolerated but showed decreased activity in both the intramacrophage and cholesterol-containing assays, especially for analog **21**. In addition, introduction of a polar methyl hydroxy substituent (**22**) was not tolerated. Collectively, these results demonstrate that the morpholine region is highly sensitive to both stereochemical and structural modifications, with the meso *cis* configuration of **1** providing the most favorable overall activity profile.

**Table 3.** *In vitro* anti-TB inhibitory activity and cytotoxicity of the morpholine modified analogs 12-20.

| Compound | R<sub>2</sub> | TB activities |  | Cytotoxicity |  |
| --- | --- | --- | --- | --- | --- |
|  |  | Intramacrophage<br>H37Rv THP-1<br>EC<sub>50</sub> (uM) | H37Rv Cholesterol<br>MIC<sub>90</sub> (uM) | HEK293T<br>CC<sub>50</sub> (uM) | HEPG2<br>CC<sub>50</sub> (uM) |
| **1** |  | 0.23 ± 0.15 | 0.26 ± 0.17 | > 20 | > 40 |
| **12** |  | 10.29 ± 10.07 | 4.55 ± 2.52 | >80 | >80 |
| **13** |  | 5.4 ± 4.04 | 9.48 ± 5.26 | 34.28 ± 8.59 | >40 |
| **14** |  | >20 | 17.45 ± 3.54 | >40 | >40 |
| **15** |  | 10.58 ± 7.66 | 0.275 ± 0.0776 | >40 | >40 |
| **16** |  | 11.61 ± 2.76 | 11.19 ± 1.25 | >40 | >40 |
| **17** |  | >20 | >20 | >40 | >40 |
| **18** |  | 18.51 ± 6.37 | 9.86 ± 9.14 | >40 | >40 |
| **19** |  | 6.67 ± 3.9 | 8.95 ± 2.55 | >80 | >80 |
| **20** |  | 0.72 ± 0.63 | 0.66 ± 0.464 | 30.04 ± 11.84 | 29.19 ± 6.27 |
| **21** |  | 1.18 ± 0.917 | 1.64 ± 1.41 | >40 | >40 |
| **22** |  | >20 | >20 | >40 | >40 |

We then turned our attention to modification of the central aromatic core of the scaffold (**Table 4**). We first examined the role of the fluorine substituent on the phenyl ring. Either removal of the fluorine (**26**) or introduction of an additional fluorine substituent (**27**) resulted in a >10-fold loss of activity, indicating a strong preference for the original substitution pattern. In contrast, replacement of the fluorine with chlorine afforded the more potent analog **25**, which exhibited an intramacrophage EC_50_ of 0.079 µM while maintaining a cholesterol MIC_90_ comparable to that of compound **1**. Replacement of the phenyl ring with heteroaromatic rings, including pyridine (**23**) and pyrimidine (**29**), resulted in a complete loss of activity. We also investigated the role of the nitrogen linker connecting the two aromatic rings. *N*-Methylation or replacement of the nitrogen atom with oxygen was detrimental to antimycobacterial activity, indicating that diarylamine motif is important for maintaining anti-TB activity.

**Table 4.** *In vitro* anti-TB inhibitory activity and cytotoxicity of the central aromatic modified analogs 23-29.

| Compound | Ar<sub>3</sub> | TB activities |  | Cytotoxicity |  |
| --- | --- | --- | --- | --- | --- |
|  |  | Intramacrophage<br>H37Rv THP-1<br>EC<sub>50</sub> (uM) | H37Rv Cholesterol<br>MIC<sub>90</sub> (uM) | HEK293T<br>CC<sub>50</sub> (uM) | HEPG2<br>CC<sub>50</sub> (uM) |
| **1** |  | >20 | >20 | >80 | >80 |
| **23** |  | >20 | >20 | >80 | >80 |
| **24** |  | >20 | >20 | >40 | >40 |
| **25** |  | 0.079 ± 0.050 | 0.27 ± 0.27 | > 40 | > 40 |
| **26** |  | 2.47 ± 0.86 | 4.36 ± 0.108 | >80 | >80 |
| **27** |  | 18.51 ± 6.37 | 7.29 ± 4.06 | 25.65 | 32.41 |
| **28** |  | >20 | >20 | >80 | >80 |
| **29** |  | >20 | >20 | >40 | >40 |

### ADME and in vivo pharmacokinetics profiling

Based on results of SAR optimization, the two most potent compounds, **1** and **25**, were selected for further evaluation of their physicochemical and ADME properties (**Table 5**). *In silico* prediction indicated that compound **25** (AlogD = 4.9) was slightly more lipophilic than compound **1** (AlogD = 4.5), consistent with replacement of the fluorine substituent with chlorine. Consistent with its higher lipophilicity, compound **25** exhibited increased plasma protein binding in both mice (99.0%) and human (98.4%) plasma compared with compound **1** (94.8% and 95.2%, respectively)

**Table 5.**
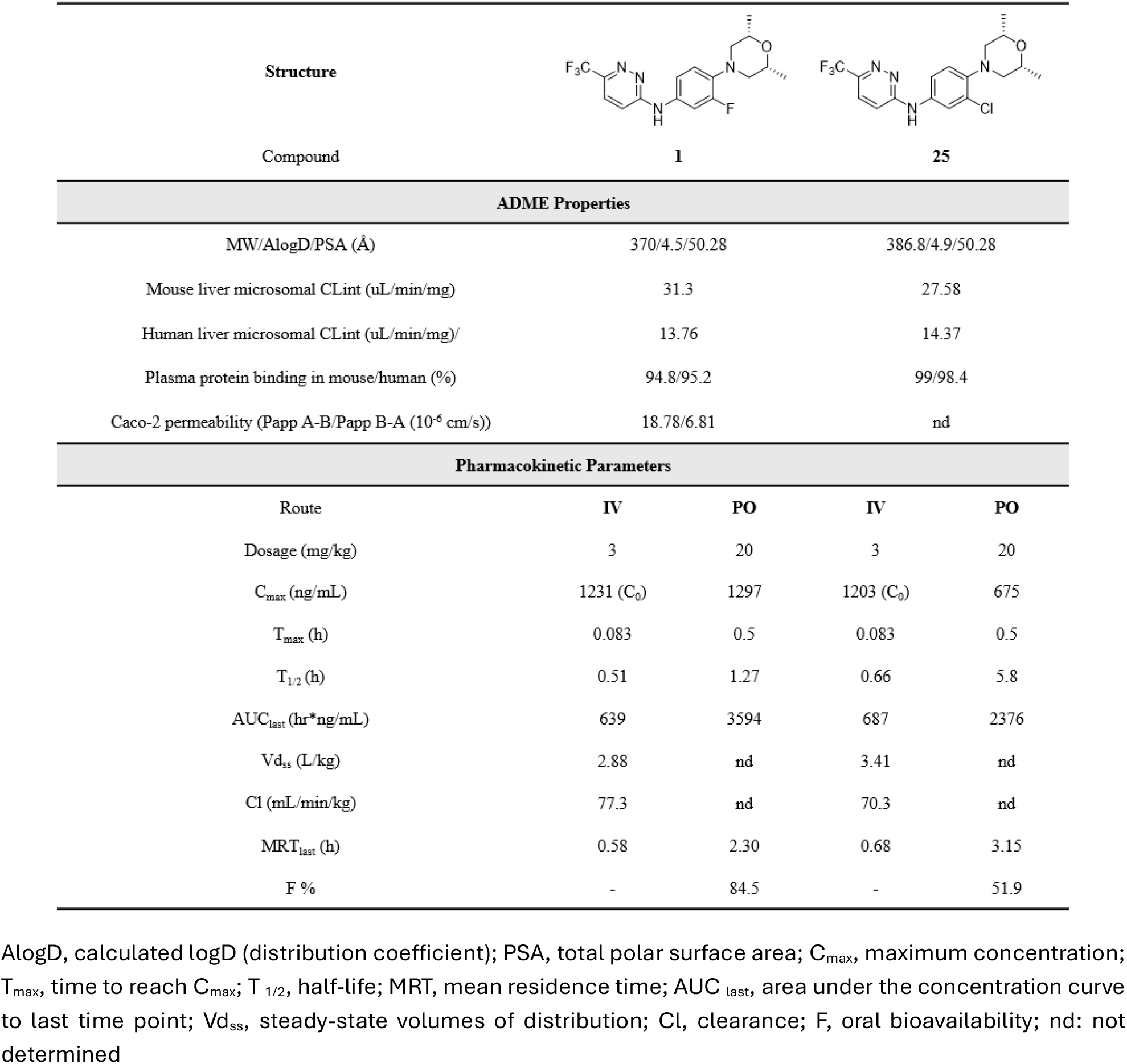
ADME and *in vivo pharmacokinetics* profiles of leading compounds 1 and 25.

Both compounds showed comparable microsomal stability, with intrinsic clearance (CL_int_) values of 31.3 and 27.58 µL/min/mg in mouse liver microsomes and 13.75 and 14.37 µL/min/mg in human liver microsomes for compounds **1** and **25**, respectively. Compound **1** also demonstrated favorable permeability in the Caco-2 assay, with no evidence of substantial efflux (efflux ratio (Paap B-A / A-B) = 0.36).

To determine oral bioavailability, compounds **1** and **25** were evaluated in mouse pharmacokinetic studies following intravenous (IV, 3 mg/kg) and oral (PO, 20 mg/kg) administration using a solution formulation of 75% PEG/25% D5W (**Table 4**). Following IV administration, compounds **1** and **25** exhibited similar pharmacokinetic profiles, with short terminal half-lives (T_1/2_ = 0.51 and 0.66 h, respectively), relatively high clearance (Cl = 77.3 and 70.3 mL/min/kg, respectively), and moderate steady-state volumes of distribution (Vd_ss_) of 2.88 and 3.41 L/kg, respectively. Following oral administration, compound **25** exhibited a substantially longer apparent terminal half-life than compound **1** (5.8 vs 1.27 h), together with a modestly longer mean residence time (MRT_last_ = 3.15 vs 2.30 h). However, compound **1** achieved an approximately 1.9-fold higher C_max_ (1297 vs 675 ng/mL) and greater systemic exposure (AUC_last_ = 3594 vs 2376 h·ng/mL). Compound **1** also demonstrated high oral bioavailability (F = 84.5%), whereas compound **25** showed moderate bioavailability (F = 51.9%).

Analysis of the plasma concentration–time profile relative to the cholesterol MIC indicated that, after a 20 mg/kg oral dose, total plasma concentrations remained above the MIC for approximately 6–8 h, suggesting that twice-daily (BID) dosing might provide more sustained exposure over MIC_90_. To further characterize the pharmacokinetic profile following repeated administration and assess the preliminary tolerability of this series, a 4-day repeat-dose study was conducted with compound **1**. Mice were administered compound **1** BID for 4 consecutive days at doses of 30, 100, or 200 mg/kg using a suspension formulation containing 0.5% methylcellulose (MC) and 0.5% Tween 80 (**Figure 3**). Plasma samples were collected on Days 1 and 4, and terminal 24h perfused lung samples were collected at the end of Days 1 and 4 for lung tissue concentration analysis. Compound **1** was well tolerated at all dose levels, with no treatment-related clinical abnormalities or body-weight loss observed. In addition, no notable findings were observed in terminal serum chemistry or hematology parameters. On day 1, both C_max_ and AUC_last_ increased in a dosed proportional manner. However, on day 4, lower-than-dose-proportional increases in C_max_ and AUC_last_ were observed, with the effect more pronounced for C_max_. Moreover, systemic exposure on Day 4 was substantially lower than that on Day 1, with AUC_last_ reduced by approximately 42%, 75%, and 82% at 30, 100, and 200 mg/kg, respectively. The magnitude of the exposure reduction increased with dose, indicating a dose- and time-dependent decrease in systemic exposure following repeated oral administration. Despite the reduced plasma exposure, lung-to-plasma concentration ratios remained above 2 at 100 and 200 mg/kg on Day 4, suggesting appreciable distribution into lung tissue. This property may be advantageous for the treatment of pulmonary TB, for which the lung represents the primary site of infection. To investigate the potential mechanism underlying the reduced systemic exposure of compound **1** following repeated dosing, nuclear receptor assays were conducted to assess activation of the human constitutive androstane receptor (hCAR) and pregnane X receptor (hPXR), both of which regulate the expression of drug-metabolizing enzymes, including CYP2B6 and CYP3A4.^19^ Compound **1** showed only modest activation of either hCAR or hPXR across the tested concentration range (0.12–30 µM; **Table S1** in supporting information) relative to the respective positive controls (phenytoin and rifampicin), suggesting that activation of these nuclear receptors is unlikely to account for the reduced exposure observed following repeated dosing. Further investigation of the underlying mechanism, together with continued SAR optimization, is ongoing to identify compounds with improved potency and pharmacokinetic profiles.

**Figure 3.**
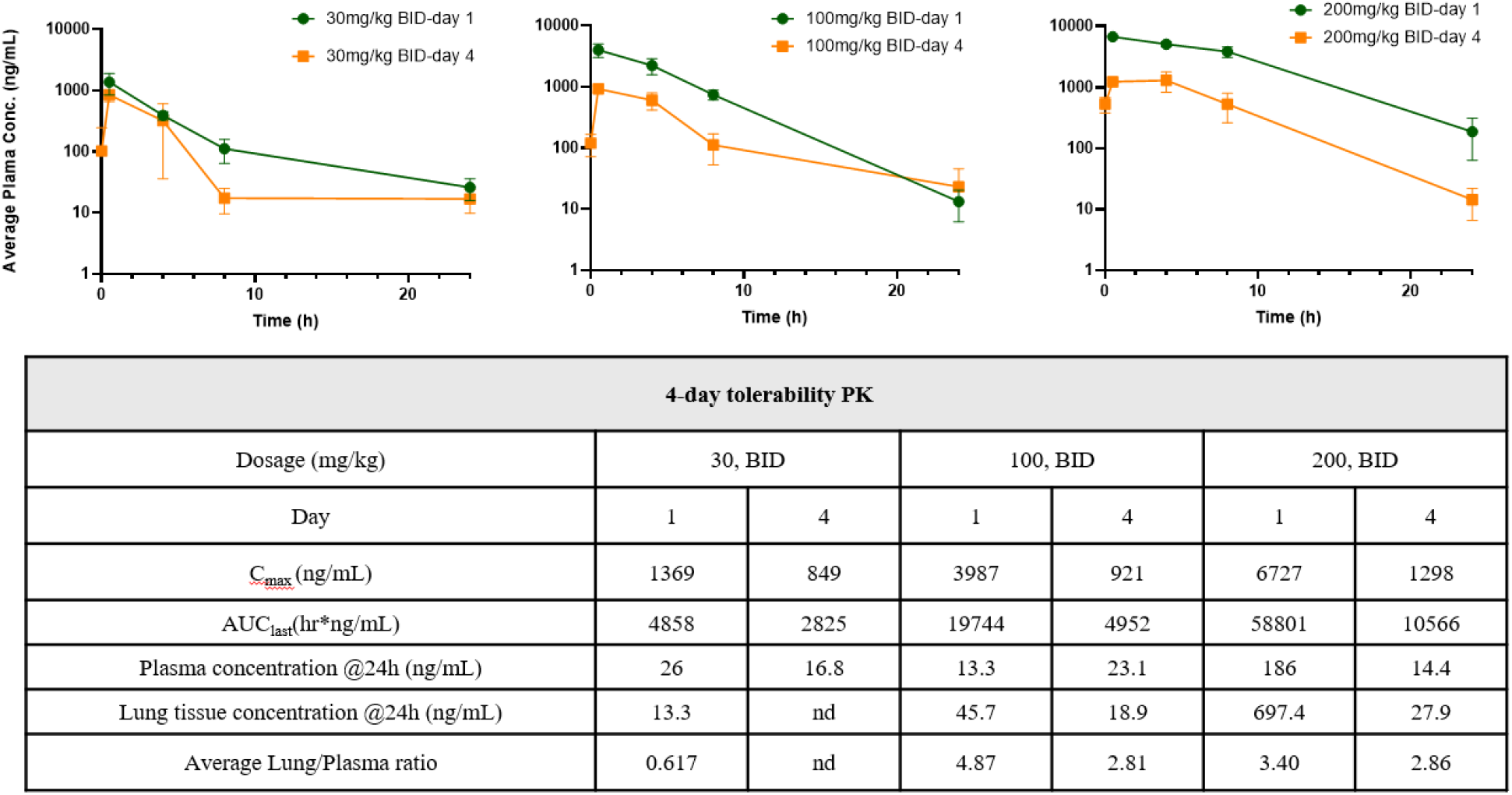
Pharmacokinetics of compound **1** following 4-day repeat dosing in mice nd: not determined

## Conclusion

We herein report the discovery, biological characterization, and SAR optimization of a novel Rv1625c agonist scaffold. Lead compounds were further evaluated for ADME and pharmacokinetic properties, and compound **1** was assessed in a 4-day repeat-dose mouse study to provide a preliminary tolerability evaluation. Although compound **1** exhibited suboptimal pharmacokinetic properties following repeated dosing, the structural novelty and promising antimycobacterial activity of this chemotype support continued optimization toward analogs with improved potency and pharmacokinetic profiles. Given the potential of Rv1625c activation as a differentiated mechanism for shortening TB treatment, this chemotype provides a valuable foundation for further optimization and supports continued exploration of Rv1625c agonists as a promising approach in TB drug discovery.

### Notes

The authors declare no competing financial interest.

## Acknowledgements

We are grateful to the Calibr compound management and pharmacology group for their assistance with this project. Special thanks to Sharon Irelan for her assistance in project management. This work was supported by a grant from the Bill & Melinda Gates Foundation #OPP1208899 to Calibr.

## Authors Contributions

The manuscript was written through contributions of all authors. All authors have given approval to the final version of the manuscript. Medicinal chemistry design and SAR optimization were performed collaboratively by the Scripps Research and Lgenia chemistry teams.

